# Dynamics of male African elephant character durability across time, age, and social contexts

**DOI:** 10.1101/2024.05.24.595367

**Authors:** Caitlin E. O’Connell-Rodwell, Jodie L. Berezin, Colleen Kinzley, Patrick T. Freeman, Monica N. Sandri, Dustin Kieschnick, Timothy C. Rodwell, Mariana Abarca, Virginia Hayssen

## Abstract

Post-dispersal male African elephants (*Loxodonta africana*) live within complex social networks. To quantify the durability of male elephant character (or personality) within these networks, we employed behavioral repeatability analysis tools across social and environmental contexts. We collected behavioral data from thirty-four individually-identified male elephants at the same waterhole over five field seasons (2007-2011) in Etosha National Park, Namibia. Using repeatability models to assess ten behavioral categories, we found five behaviors (affiliation, aggression, dominance, self-directed anxious, and self-directed comfort) were consistent at the individual level. Interestingly, some of these behaviors were also significantly repeatable, depending on social context. In particular, the presence of younger males and a keystone male (i.e., the most dominant and socially-integrated individual during our study period) had the biggest impact on adult male behaviors. Surprisingly, the presence of elephants in musth had little impact. Finally, we found that younger individuals were more alike in their overall character profiles than older males, further supporting the hypothesis that male elephants develop unique, yet socially-flexible character types as they age. These results demonstrate that male elephants possess distinct character traits that are also behaviorally adaptable, depending on the social context. Overall, our research further uncovers the complexity of male elephant individuality and social dynamics that can be leveraged to improve in-situ and ex-situ management and conservation decisions for the species.

## Introduction

Individual social animals within a population often express certain behaviors—or sets of correlated behaviors—differentially, and may do so consistently or flexibly across time, space, and environmental gradients [1–4]. More recently, the field of Conservation Behavior has emerged in an effort to understand which behaviors have important implications for the conservation of vulnerable species, as well as the mechanisms that enable some animals to succeed or fail to adapt in a rapidly changing world [5]. However, given the ubiquity and diversity of behaviors present across taxa, scientists have the challenging task of determining which species, demographic groups (e.g., juveniles or adults), behavioral traits (e.g., aggression or affiliation), or study contexts should be prioritized [5]. Likewise, researchers must also decide how behaviors should be measured in order to best inform wildlife management and conservation policies.

The African savannah elephant (*Loxodonta africana*) is a highly intelligent, socially complex, and long-lived megafaunal species that has been a conservation priority over the last several decades. Like other large mammalian herbivores, elephants provide key ecosystem services [6] and are both culturally and economically significant, yet are severely impacted by anthropogenic disturbance and climate change [7]. One way to improve conservation efforts is through behavioral repeatability research, repeatability of behavior being a proxy for animal character (often referred to as personality, temperament, or consistent individual differences) at the population level [1,2].

To date, all three species of elephants are known to display behavioral repeatability. However, the current body of research on elephant character is based largely on captive [8–15] and semi-captive elephants [16–19], with male data being grouped with female data, and only two studies on wild elephants to date (*L. cyclotis* - [20]; *L. africana* - [21]). While these previous studies provide the foundation for elephant character research, their results may not generalize well to free-ranging systems, given the inconsistencies between these two environmental contexts.

In addition, research on elephant character and behavioral repeatability have been primarily focused on females. Unlike female elephants who spend their entire lives in family groups, males traverse dynamic all-male societies and spend more time alone than their female counterparts due to sex-based differences in reproductive strategies [22].

Male elephant society is dynamic, consisting of dominance hierarchies [23], complex social networks and associations [23–26] based on long-term relationships [26], kinship [24], and age structure [25,27,28]. These differences in life history likely impact the expression of consistent behaviors. The lack of research on male elephants is likely due to the limited sample sizes in captivity, as well as the challenge of collecting long-term, individual-based behavioral observations for a highly mobile species in the wild.

The goal of this study was to determine whether free-ranging male elephants of post-dispersal age differentially express behaviors consistently and/or flexibly as a function of time, age class, and social context. Using a rich collection of long-term behavioral data from individually-identified male elephants in Etosha National Park, Namibia, we first quantified the repeatability of ten major behavioral categories in order to establish the most stable character traits in our study system. Our second aim was to determine which social contexts (i.e., presence of younger males, males in musth, or a keystone individual) explain the variation observed in character traits. Finally, our third aim was to test for age-based similarities and differences in male elephant character profiles to uncover possible developmental patterns and effects of group cohesion on elephant behavior.

## Methods

### Study site

As part of a long-term elephant monitoring project that started in 1992, behavioral observations were collected on male elephants at the Mushara waterhole (hereafter Mushara) in the northeastern corner of Etosha National Park (ENP), Namibia from 2005-2011. We chose a 5-year subset of this data to analyze, collected from 2007-2011, during week field seasons from mid-June to the end of July for each consecutive year. This subset of data contained the most consistent information on the selected study subjects relevant to this research question. ENP is a fenced park that encompasses 22,970 km^2^ [29] and supports an elephant population of approximately 2,400 individuals [30]. The Mushara waterhole is fed by a permanent, artisanal spring and is the only source of drinking water within a 10 km radius [31]. The waterhole is situated within a 0.22 km^2^ clearing [32].

Behavioral observations were collected from an 8-meter-tall research tower situated about 80 meters north of the waterhole with a 360-degree view of the clearing. Elephants enter the clearing from eight well-traveled paths from the brush and usually walk directly to the water trough or pan (see [32,33]). Water flows from the source of the spring into a trough, the head of which has the freshest water and is the preferred location to drink.

### Elephant identification and age classification

Many elephants that visit Mushara have been individually identified and tracked across years using morphological traits such as ear-tear patterns, tail-hair configuration, tusk size and shape, and body size [23,34]. Male elephants are assigned a relative age class based on shoulder height, hind foot length, and facial characteristics [35]. Age classes include: one-quarter (1Q), 10-14 years old; two-quarter (2Q), 15-24 years old; three-quarter (3Q), 25-34 years old; full, 35-49 years old; and elder, 50 years and older.

A total of 34, non-musth males were included as the focal subjects in this study, while the presence of musth males was considered a social context (see section 2.5 for more details). We chose individuals who were observed at least three times across two years (mean occurrence per individual = 15.4 ± 10.0, range = 4, 40), with a mean of 21 individuals observed per year and a range of 14 to 28 individuals (S1 Table). Five males changed age classes during their years of observation [35]. Since age classes for males span approximately 10 years, we did not anticipate abrupt shifts in the expression of behaviors observed in these five individuals. As such, these individuals were categorized by the age class they remained within for most of the study (i.e., 4 out of 5 years). For the 34 males, four were classified as 1Q, six were 2Q, nine were 3Q, twelve were full, and three were elders.

### Behavioral observations

Elephant behavioral observations were recorded from approximately 11:00 am to 5:00 pm when elephants are visible and easily distinguishable. Behavioral data were collected as “events”, where events began when one or more elephants entered the clearing and ended when all elephants left the clearing. If an elephant was alone, the event was terminated after fifteen minutes. Since the focus was on male elephants, events with female elephants were removed. In cases where females arrived during an event, we kept the part the event before the females arrived. Behavioral observations were collected using all-occurrence sampling [36] for all elephants present at the waterhole using a customized datalogger programmed with Noldus Observer software (Noldus Information Technology Inc., Virginia, USA).

Sixty-six distinct behaviors were recorded (Table 1; S2 Table). To include the full suite of unique behaviors that male elephants display and to avoid removing low-occurrence behaviors, we pooled behaviors with a similar context (or expressions of similar behavioral patterns) into ten categories [23,37]. The fine-scale behaviors in each category and category definitions are listed in Table 1, while definitions for the fine-scale behaviors are provided in S2 Table. Moving forward, these behavioral categories will be referred to simply as “behaviors.” The final ten behaviors include affiliation, aggression, displacement, escalated aggression, play, retreat, self-directed anxious, self-directed comfort, social contentment, and vigilance.

**Table 1.** Condensed ethogram describing the ten behavioral categories and list of sixty-six discrete behaviors that make up the behavioral categories. Behaviors and categories are adapted and modified from [23,37]. For behavior categories, n represents the total occurrence of discrete behaviors within that category, pooled across individuals, events, and years.

| **Behavior (category)** | **n** | **Behavior category definition** | **Fine-scale behaviors** |
| --- | --- | --- | --- |
| Affiliation | 2,461 | Positive interaction between individuals; aids in relationship building and maintenance. | Backs into; Body to body; Ear on face; Ear on rear; Follows; Foot to body; Head to body; Head to head; Inspecting; Mount; Pre-mount; Pushes; Rubs; Tail to body; Trunk to body; Trunk to head; Trunk to mouth; Trunk to temporal; Trunk to tusk; Trunk wrap |
| Aggression | 2,038 | Negative interaction to intimidate or threaten other elephants; no physical contact made between elephants. | Aggressive ear flap; Ear fold; Ears held out; Foot toss; Hard ear flap; Head held up; Head shake; Open mouth threat; Tail slap; Trunk drag; Trunk fist; Trunk throw |
| Dominance | 872 | Displacement of another individual; used to calculate the dominance hierarchy. | Displacement |
| Escalated aggression | 69 | Negative interaction to threaten or attach other elephants; often no physical contact. | Charge; Chase; Combat; Head down; Head down; Head thrust; Lunge; Pursue; Stand off |
| Play | 493 | Social play where individuals engage in exaggerated or loose movements and friendly sparring that avoids injury. | Gentle sparring; trunk swing |
| Retreat | 206 | Avoidance of aggression from another individual. | Back up; Retreat |
| Self-directed anxious | 1,882 | Nervous and reactive behaviors directed towards oneself; often to attend to the actions of other elephants or after receiving aggression from another. | Foot rubs; Tail lift; Touches own tusk; Trunk own mouth; Trunk own temporal; Trunk suck; Trunk twist |
| Self-directed comfort | 498 | Comfort or relaxed behaviors directed towards oneself. | Cross leg; Rest trunk; tusk hang |
| Social Contentment | 1,992 | Group activity; relaxed behaviors observed during times of stillness at the waterhole. | Ear flap; Tail swing |
| Vigilance | 4,616 | Environmental assessment, often towards a conspecific to attend to the actions of other elephants; vigilance behaviors directed toward human activity were removed. | Freeze; Freeze trunk ground; Look; Orient; Over shoulder; Smell |

### Identifying the keystone individual

We wanted to examine the degree to which the presence of key social actors within the population may affect behavioral repeatability. For this social context, we identified a keystone individual based on two facets of social importance: 1) social network centrality and 2) dominance rank. Prior to network and dominance analyses, we filtered the dataset to (1) remove all individuals without a positive identification (i.e. unknown individuals), (2) remove all individuals recorded as being in musth during the observation year, and (3) remove all positively-identified individuals that were recorded at fewer than three independent observation sessions (events) during the season.

*1) Social network centrality:* For each year, we constructed association networks based on co-presence at the waterhole during observation sessions. For the purposes of our analyses, we applied the Gambit-of-the-Group approach [38], assuming that all individuals recorded during the same observation session were associating with each other, even if they arrived and/or departed at different times.

For each year, we built weighted matrices of dyad-level association indices based on the Simple Ratio Index of association (SRI; [39,40]) with larger association indices indicating individuals were more closely associated, and from there, calculated individual eigenvector centrality scores. This network metric is frequently used to quantify an individual’s influence on the broader network by assessing both its direct connections and the connections of its neighbors. An animal is considered more central if it is connected to other individuals that are also well-connected in the social network [41]. This metric helps identify key individuals that may play influential roles in information flow or social dynamics within the animal group. Networks and association indices were calculated using the asnipe package [42] and eigenvector centrality was calculated using the igraph package [43].

1. *2) Social dominance hierarchy:* In this study system, the displacement of an individual—defined as an instance where one elephant forces another to change his position and move away, possibly so that the initiator can take the position [23]—is an obvious non-combat behavior used to express dominance. We used dyad-level displacement contests to construct an ordinal dominance hierarchy per year and identify the most dominant individual in the population.

We calculated ordinal dominance hierarchies using the normalized David’s Score (DS). DS estimates an individual’s dominance by considering the proportion of an individual’s dominance interactions result in wins or losses across all the dyads with whom he interacts while considering the total number of dominance interactions observed. The highest values are assigned to individuals that most consistently win their contests [44–46]. Raw DS are converted to normalized David’s scores, such that in a population of N individuals, scores vary between 0 and N-1, with the highest value identifying the most dominant individual in the defined population (see [45] for derivation). Displacement contest matrices were constructed using the *Perc* package [47] and normalized DS were calculated using the *EloRating* package [48].

### Social context descriptions

We assessed behavioral flexibility across different social contexts at the waterhole. The waterhole acts as both an important resource and a place where a large suite of social interactions among elephants can be easily observed. We chose three social contexts that occurred frequently at Mushara waterhole and that we hypothesized, based on long-term observations, appear to influence male behavior: the presence of a musth male, the presence of a keystone male, or the presence of young males. We hypothesized that these social contexts would illicit behavioral flexibility in varying degrees for each behavior.

In total, eight social contexts were defined in the study (Table 2). The presence of musth males, the keystone male, and young males (defined as individuals in the 1Q and 2Q age classes) were recorded for each event. We used presence-absence coding to then categorize events into the three social contexts. To account for events with an overlap of social contexts (e.g., musth and young males present), an additional four contexts were added. The final category was events with only adult, non-musth males (age classes 3Q, full, and elder). This category was considered the baseline social context and labeled as “adults only.” Due to the low occurrence of events where both a musth male and the keystone male were present (n = 1), as well as the musth male, keystone male, and young males were present (n = 2), these intervals were removed from subsequent analyses, leaving 6 total social contexts.

**Table 2.** Social context categories, descriptions, and the total interval occurrences. Young males are those in the 1Q and 2Q age classes and older males (“adults”) are those in the 3Q, full, and elder age classes. The social contexts of musth male and keystone male (n = 1), and musth male, keystone male, and young males (n = 2) are not displayed here due to low sample sizes.

| Social context category | Description | Total Occurrences |
| --- | --- | --- |
| Musth | Musth male(s) present with other males in the older age classes. The keystone and young males are not present. | 21 |
| Keystone | The keystone male is present with other males in the older age classes. Musth and young males are not present. | 13 |
| Young | Young males present with or without older males. The keystone male and musth males are not present. | 55 |
| Musth and young | Musth and young males present with or without older males. The keystone male is not present. | 14 |
| Keystone and young | Keystone male and young males are present, with or without older males. Musth males are not present. | 26 |
| Adults only | Only older males (3Q+) are present. Musth, keystone, and young males are not present. | 68 |

### Intervals for changing social contexts

The social context often changed during long events. To account for this, an interval was assigned each time the social context changed within an event. As an example, some adults and young males were present at the start of the event and the interval was labeled as “1.” Then, the keystone male arrived, so the first interval of the event ended, and the second interval began. Intervals with solitary elephants, family groups, or only unknown males were removed. Across the five years, this left a total of 148 events (mean per year ± SD: 29.6 ± 10.41) that contained 200 intervals (40.0 ± 13.96) (S1 Table). The mean time per event was 70.14 ± 47.10 minutes (range = 6.95, 249.39) and the average time per interval was 49.61 ± 33.76 minutes (range = 5.19, 178.72).

### Statistical Analysis

All statistical analyses were conducted using R (version 4.3.1)[49]. Significance was evaluated at α = 0.05 for all models.

### Repeatability analysis

We used repeatability models to assess the consistency of elephant behaviors. Repeatability (*R*) is an intra-class correlation measure that is used in animal personality research to quantify stable individual differences in behavior at the population level [50]. Repeatability models use the mixed-effects model framework, where the random effect estimates variance of repeated behavior measures of individuals. *R* is calculated as the group-level variance over the sum of the group-level and residual variances. In other words, where low intra-individual variation and high inter-individual variation creates a high repeatability, or *R* value [2].

Behavior rates were calculated per year per elephant as the frequency of each behavior within an event over the total time the individual was present in that event. Rates were then transformed using the natural log due to significant right skew for each of the ten behaviors. Individuals that had less than two rates for a given behavior were removed from the model.

We fit a total of 10 models using the ‘rpt’ function in the *rptR* package (Stoffel et al. 2017). The models were fit with the ln-transformed behavior rate as the response variable, and with crossed, random effects of year, event, and individual ID. Year and event were included as random effects to account for sampling structure. Additionally, we calculated the repeatabilities for year and event to evaluate the impact that time (year) and social context (event) has on behavior rates, where significant repeatabilities indicate that behavior rates were stable within events/years and differ amongst events/years. Each model was specified with a Gaussian distribution with 1000 bootstraps for the calculation of confidence intervals and *p*-values. All model residuals were inspected to confirm assumptions of homoscedasticity and normality of residuals were met.

The rptR package calculates confidence intervals using parametric bootstrapping and *p*-values using likelihood ratio testing, leading to conflicting results such as significant p-values and confidence intervals containing zero [51,52]. In addition, repeatability values reported in nonhuman animals are typically low [2]. As such, we determined whether behaviors were repeatable using a combination of the *R* value, confidence intervals, and *p-* values [52]. We considered behaviors repeatable if the *R* value was greater than 0.1, did not contain 0 in the confidence interval, and had a significant *p*-value.

### Behavioral flexibility among social contexts

We used linear mixed-effects models to assess the impact of social context on behaviors that had significant event repeatability. For this analysis, we focused only on the rates observed for adults (age classes 3Q, full, and elder) and removed the rates for young males (age classes 1Q and 2Q) and the keystone male. This method allowed for a direct comparison of how adults change their behavior rates in social contexts and removed the impact that the keystone male or young males have on behavior rates. For this analysis, we re-calculated behavior rates at the interval level (see section 2.6), rather than the event level (as was done for the repeatability analysis) to measure the impact that social contexts had on adult male elephant behavioral consistency.

We fit a model for each of the behaviors with a high event repeatability, with behavior rate as the response variable and social context as the fixed effect. Nested random-effects of year, event, and interval were included, as well as a partially-crossed random effect of individual ID to account for repeated measures of individuals. Behavior rates were first transformed using the natural log to meet normality assumptions. Models were fit using the ‘lmer’ function in the *lmerTest* R package [53]. This package builds on the lme4 package by including denominator degrees of freedom and *p*-values estimated by Satterthwaite’s method. Estimated marginal means were calculated for each model using the ‘ggpredict’ function in the *ggeffects* R package [54]. Model assumptions of linearity, homoscedasticity, and normality were assessed using functions from the R package *DHARMa* [55]; assumptions were met for all models.

### Age related differences in character profiles

To assess the similarities of the character profiles among individuals, we used a non-metric multidimensional scaling (NMDS) clustering analysis. We used NMDS clustering since the behavior rate data was not suitable for traditional, personality data-reduction techniques (e.g., factor or principal components analysis) due to the correlation matrix failing to meet the Kaiser-Meyer-Olkin (KMO) and Bartlett’s test of sphericity criterion [56]. NMDS clustering is typically used for assessing similarities in species compositions among communities, where the number of individuals of each species is recorded per site [57]. An important assumption is that sampling effort is uniform across sites, so the abundances of species between sites are directly comparable. Due to differences in occurrence (total time each individual was observed) among individuals, count data were not appropriate, and proportions were required. For each individual, we calculated the proportion they displayed each repeatable-by-individual behavior, which we’ve termed an individual’s character profile.

Since proportions are compositional data, characterized as being bound by a lower and upper limit and existing in ‘simplex’ space, the data need to be transformed to be brought into ‘real’ space using the Aitchison’s distance [58]. Aitchison’s distance is calculated by transforming compositional data using the centered log-ratio (clr) and then calculating the Euclidean distance [59,60]. A modified version of the clr transformation was used, called the robust clr [61] to account for individuals who do not display one of the behaviors (n = 7). The robust clr allows for “true zeros” to remain in the dataset without affecting downstream analyses or removing the individuals who did not display a behavior [61]. We considered the lack of these behaviors as a true zero, since the elephants did not display these behaviors during the observation time. As such, data transformations were performed on all nonzero values. The Euclidean distance matrix was then calculated with the robust clr transformed data using the ‘vegdist’ function in the R package *vegan* [62], specified with pairwise deletions to retain the zeros.

We used NMDS to express variation in two-dimensional space and validated the analysis by calculating the stress value (values smaller than 0.2 are acceptable; [63]). Model fit was further evaluated using two methods: a goodness of fit plots of individuals, and a Shepard diagram of the linear and non-metric fit of the observed dissimilarities and their relation to the ordination distance.

The resultant NMDS cluster plot was overlaid with the five elephant age classes to visually assess the relationship between age and relative position of individual character profiles. We used an Analysis of Similarities (ANOSIM) model to statistically test the difference between the five age classes. We used the ‘anosim’ function in the R package *vegan* [62] to fit each model with the dissimilarity matrix calculated for the NMDS model. The test statistic, *R*, is scaled from 1 to −1, where values greater than 0 means objects are more dissimilar between groups than within groups, and the opposite for values below 0 [63]. Groups were considered significantly different from each other if the *R* value was greater than 0 and the corresponding *p*-value was significant.

## Results

### Social network centrality and dominance hierarchy analyses

The goal of our network centrality analysis was to determine whether a single full-size adult male maintained the highest network eigenvector centrality across all five years. The number of individuals included in the annual networks varied substantially after applying filtering criteria (mean = 25, range = 8, 27 individuals; Figure 1a). Our network analyses indicated that one individual, a full-sized adult male (Male #22) had the highest average eigenvector centrality of all individuals included in our analysis across all five years (mean = 0.91; SD =0.18). This male was also the only full-size adult that was consistently sighted (i.e. observed at three or more independent observation events during a season) across all five years of the study.

**Figure 1.**
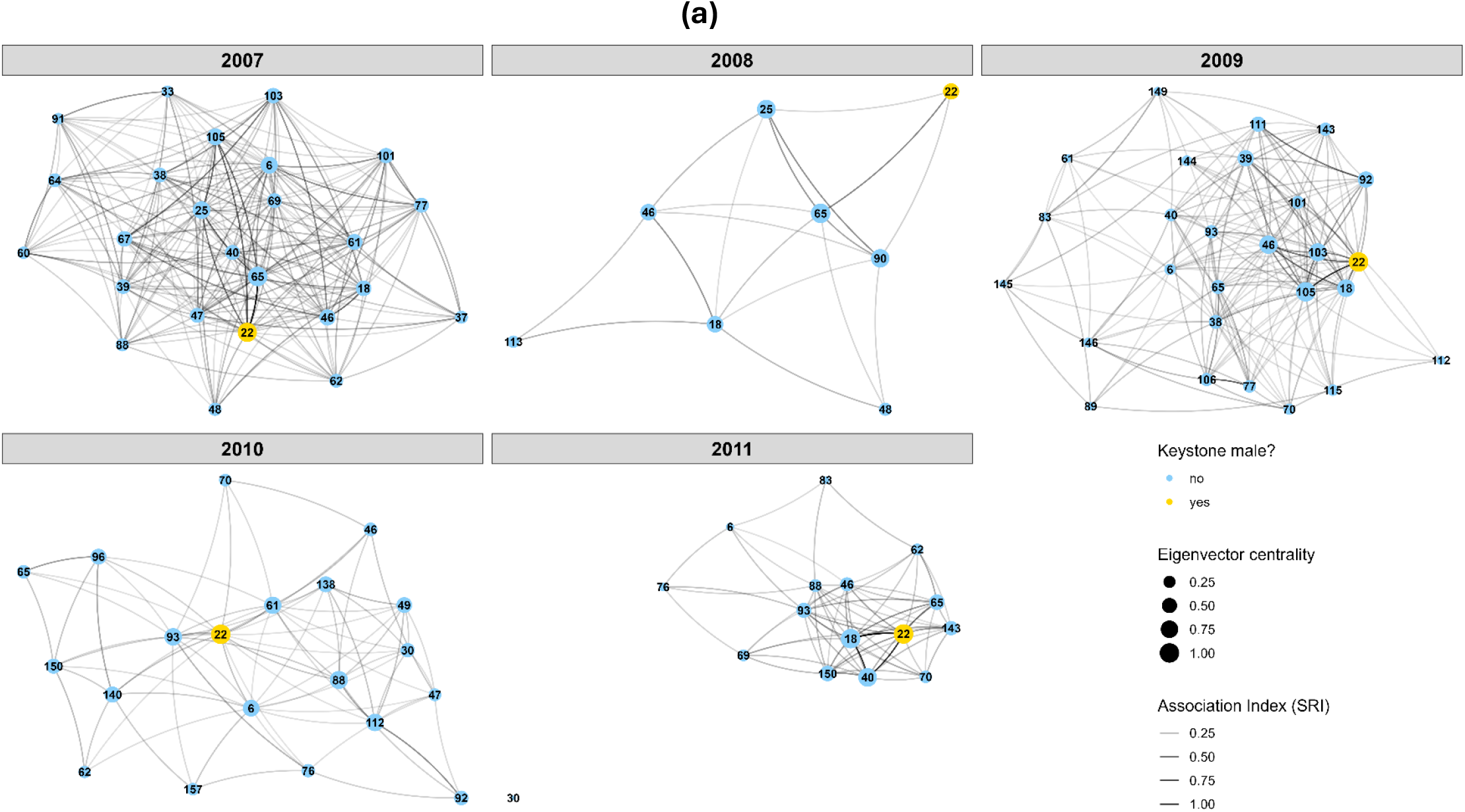
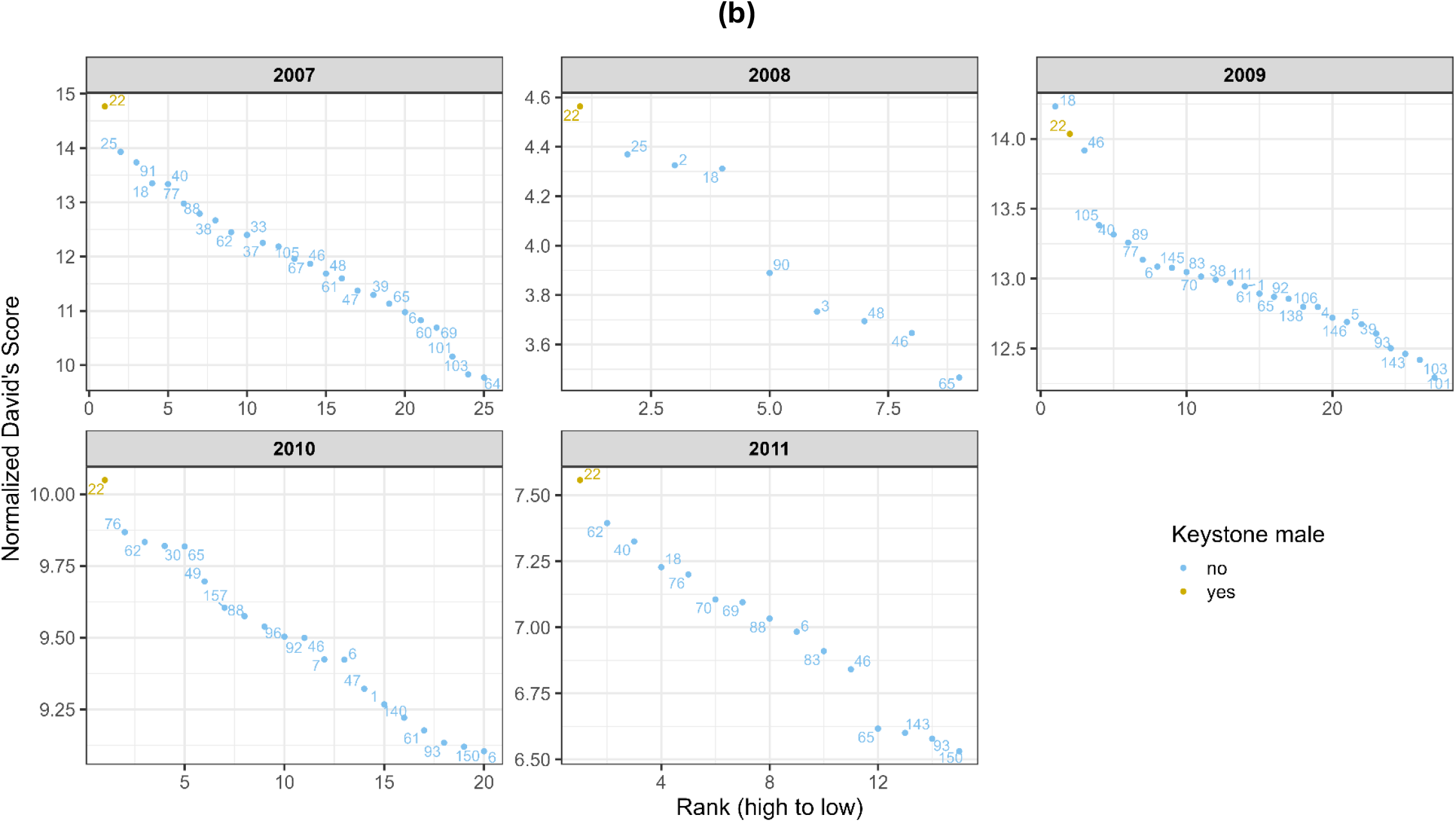
Social network and dominance hierarchy for each of the five years (2007-2011). The keystone individual (#22) is highlighted in yellow. (a) Social network centrality with a varied number of individuals per year (mean = 25, range = 8, 27). Circles represent individuals and the size of the circle represents how central they are to the network, binned into four levels. The lines between the individuals represent their association index relative to the individual the line is connecting to, where darker lines represent strong associations (also binned into four levels). (b) Dominance hierarchy with a varied number of individuals per year (mean = 25, range = 9, 27). Each point and corresponding number represent an individual. Due to differences in the number of individuals and David’s Scores in each year, both the x and y axes are displayed on different scales to better display the rankings.

In addition to determining whether a single adult male maintained a central place in the broader male social network, we also wanted to determine whether the same male maintained a consistently high dominance rank across the years in our study. Our displacement-based dominance hierarchy analyses suggested that this was indeed the case. Annual hierarchies were composed of an average of 25 males of all age classes (range = 9, 27). The same individual identified as consistently having the highest eigenvector centrality (Male #22) also had the highest normalized David’s Score in all but one of the study years (2009) in which he had the second highest score, but with a very small difference between him and the highest-ranking adult male of 0.2 (Figure 1b).

Taken together, our analyses suggest that the same individual was consistently the most dominant and most central to the network of male elephants over the study period. With this established, we hypothesized that this individual was likely to have a significant impact on the behavioral patterns of other males and could be considered a keystone individual.

### Behavioral repeatability

We tested whether any of the ten behaviors were repeatable across time and social context at the individual level. Five of the ten behaviors were significantly repeatable-by-individual (Table 3). All repeatabilities were low except self-directed comfort which had the highest repeatability estimate ± SE (*R* = 0.42 ± 0.10). Affiliation (*R* = 0.17 ± 0.05) and dominance (R = 0.17 ± 0.06) had the next highest repeatabilities, followed by aggression (*R* = 0.14 ± 0.05) and self-directed anxious (*R* = 0.14 ± 0.05).

**Table 3.** Repeatability estimates (individual, year (time), event (social context)) calculated for each behavior. Sample sizes (n) are presented as the number of individual elephants (number of behavioral rates). SE = standard error; CI = confidence interval. Significant repeatabilities are bolded.

| Behavior | n | Individual Repeatability |  |  | Year Repeatability |  |  | Event Repeatability |  |  |
| --- | --- | --- | --- | --- | --- | --- | --- | --- | --- | --- |
| | | $R \pm SE$ | 95% CI | $p$ | $R \pm SE$ | 95% CI | $p$ | $R \pm SE$ | 95% CI | $p$ |
| Affiliation | 34(408) | **0.17 ± 0.05** | **0.07, 0.28** | **< 0.001** | 0.02 ± 0.03 | 0, 0.09 | 0.264 | **0.21 ± 0.05** | **0.11, 0.31** | **< 0.001** |
| Aggression | 34(429) | **0.14 ± 0.05** | **0.06, 0.23** | **< 0.001** | 0.04 ± 0.04 | 0, 0.14 | 0.012 | **0.27 ± 0.05** | **0.17, 0.38** | **< 0.001** |
| Dominance | 30(310) | **0.17 ± 0.06** | **0.05, 0.30** | **< 0.001** | 0.01 ± 0.02 | 0, 0.06 | 0.245 | **0.15 ± 0.06** | **0.03, 0.27** | **0.006** |
| Escalated aggression | 16(52) | 0 ± 0.07 | 0, 0.23 | 1 | 0 ± 0.06 | 0, 0.22 | 1 | 0.36 ± 0.23 | 0, 0.75 | 0.082 |
| Play | 26(154) | 0.10 ± 0.06 | 0, 0.23 | 0.009 | 0.03 ± 0.05 | 0, 0.16 | 0.29 | **0.34 ± 0.10** | **0.12, 0.52** | **0.003** |
| Retreat | 21(106) | 0.01 ± 0.04 | 0, 0.13 | 0.412 | 0 ± 0.04 | 0, 0.13 | 1 | **0.50 ± 0.13** | **0.20, 0.7** | **0.006** |
| Self-directed anxious | 34(414) | **0.14 ± 0.05** | **0.05, 0.23** | **< 0.001** | 0.08 ± 0.07 | 0, 0.24 | < 0.001 | **0.28 ± 0.06** | **0.18, 0.4** | **< 0.001** |
| Self-directed comfort | 24(154) | **0.42 ± 0.10** | **0.20, 0.60** | **< 0.001** | 0 ± 0 | 0, 0 | 1 | 0.03 ± 0.05 | 0, 0.18 | 0.355 |
| Social contentment | 32(362) | 0.07 ± 0.03 | 0.01, 0.14 | < 0.001 | 0 ± 0.2 | 0, 0.05 | 1 | **0.46 ± 0.06** | **0.34, 0.57** | **< 0.001** |
| Vigilance | 34(502) | 0.05 ± 0.03 | 0.01, 0.11 | < 0.001 | **0.12 ± 0.08** | **0.001, 0.29** | **< 0.001** | **0.31 ± 0.06** | **0.21, 0.42** | **< 0.001** |

Only vigilance behaviors showed a significant effect of year (R = 0.12 ± 0.08). For four of the five repeatable-by-individual behaviors (affiliation, aggression, dominance, self-directed anxious), event repeatabilities were significant and similar to, or higher than, the individual repeatabilities (Table 3). Four behaviors that were not significantly repeatable-by-individual were significantly repeatable-by-event: retreat had the highest event repeatability (*R* = 0.50 ± 0.13), followed by social contentment (*R* = 0.46 ± 0.06), play (*R* = 0.34 ± 0.10), and vigilance (*R* = 0.31 ± 0.06). Escalated aggression was not significantly repeatable for any of the random effects. However, the event repeatability was high at *R* = 0.36 ± 0.23, suggesting some impact of social context.

### Behavioral flexibility among social contexts

Due to the significant event repeatabilities for eight of the ten behaviors, we expected social context to have an impact on the behavioral rates for adult male elephants (age classes 3Q, full, and elder). The presence of the keystone male and young males (age classes 1Q and 2Q) had the most impact on adult male elephants across behaviors (Figure 2; S3 Table). When the keystone male was present, the rate of affiliation behaviors significantly decreased (estimate = −0.72, SE = 0.27, p = 0.009). When young males were present, the rate of affiliation (estimate = 0.41, SE = 0.18, p = 0.021) and dominance (estimate = 0.58, SE = 0.16, p < 0.001) behaviors significantly increased, while the rate of vigilance behaviors decreased significantly (estimate = −0.31, SE = 0.13, p = 0.021). When both the keystone male and young males were present, significant decreases in aggression (estimate = −0.46, SE = 0.18, p = 0.014), self-directed anxious (estimate = −0.50, SE = 0.16, p = 0.003), social contentment (estimate = −0.73, SE = 0.22, p = 0.001), and vigilance (estimate = −0.70, SE = 0.16, p < 0.001) were observed. Finally, the rate of retreat behaviors increased slightly significantly when musth and young males were present (estimate = 0.82, SE = 0.41, p = 0.050).

**Figure 2.**
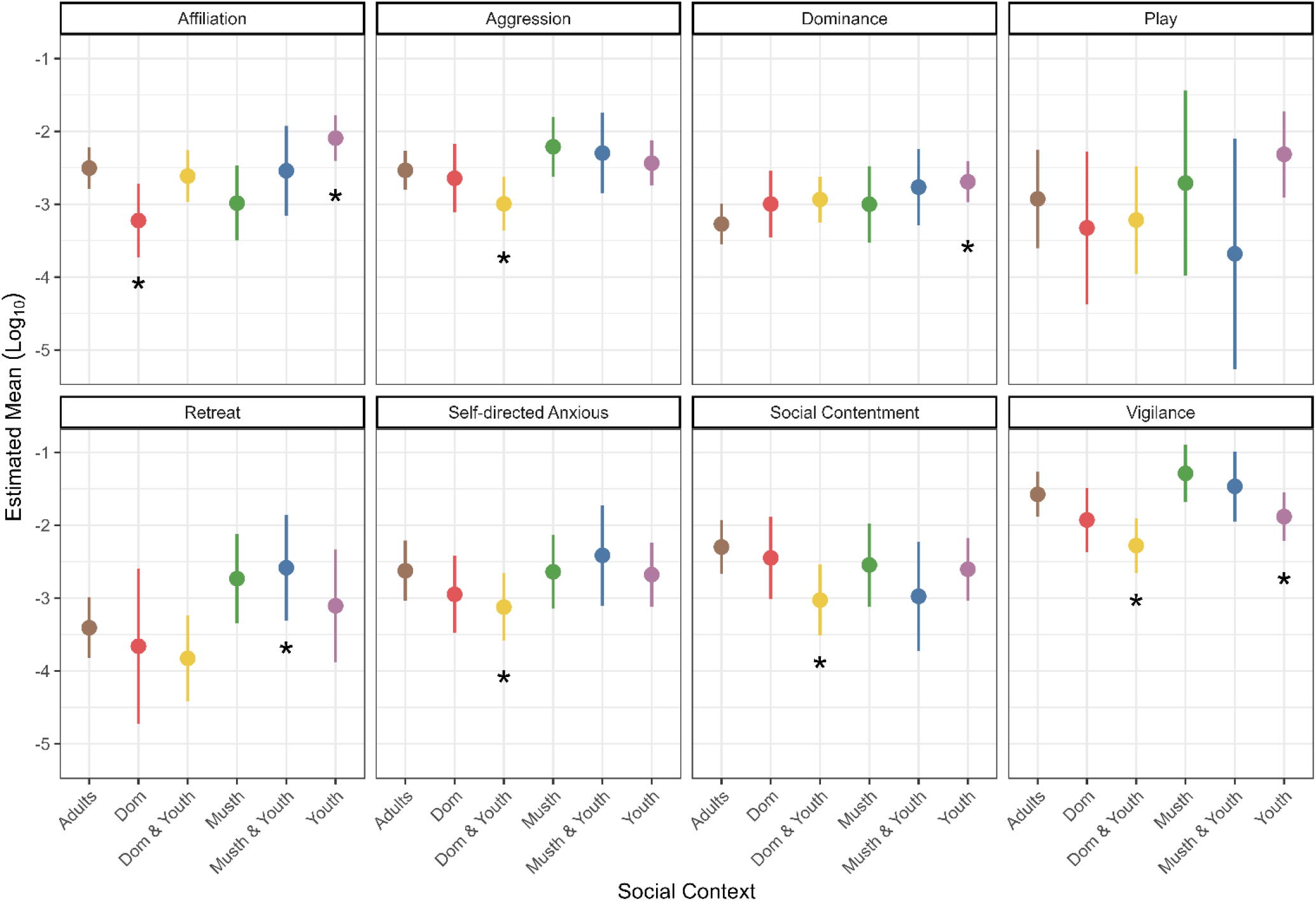
Marginal means for each of the eight behaviors with high event repeatability in six different social contexts. Only rates of adults (age classes 3Q, full, and elder) are included in this analysis. The social context refers to the presence of the keystone male, musth males, and young males, but does not include their behavior rates. The mean rates are displayed using the natural log transformation. Sample sizes and details of models are reported in S3 Table. Error bars represent the upper and lower 95% confidence intervals. Asterisks represent contexts that differ significantly from the “adults only” baseline category.

### Age related differences in character profiles

We assessed whether character profiles, an individual’s unique make-up of the repeatable-by-individual behaviors, were related to age class. NMDS analysis revealed that many of the 34 individuals display similar character profiles, while some individuals are dissimilar to most of their conspecifics (Figure 3, stress value of 0.08). The ANOSIM model indicated significant differences between the five age classes (*R* = 0.176, *p* = 0.008). 1Q, 2Q, and full-size males grouped closely within age class and occupied nearly separate areas on the figure, with the exception of one full size male grouping with the 2Q’s. Elder males grouped closely together and occupied a small area within the full-size male ellipse, suggesting similarities in character profiles among the elders. 3Q males were spread across the range of all other age classes, suggesting they display a variety of character profiles with similar attributes to the four age classes.

**Figure 3.**
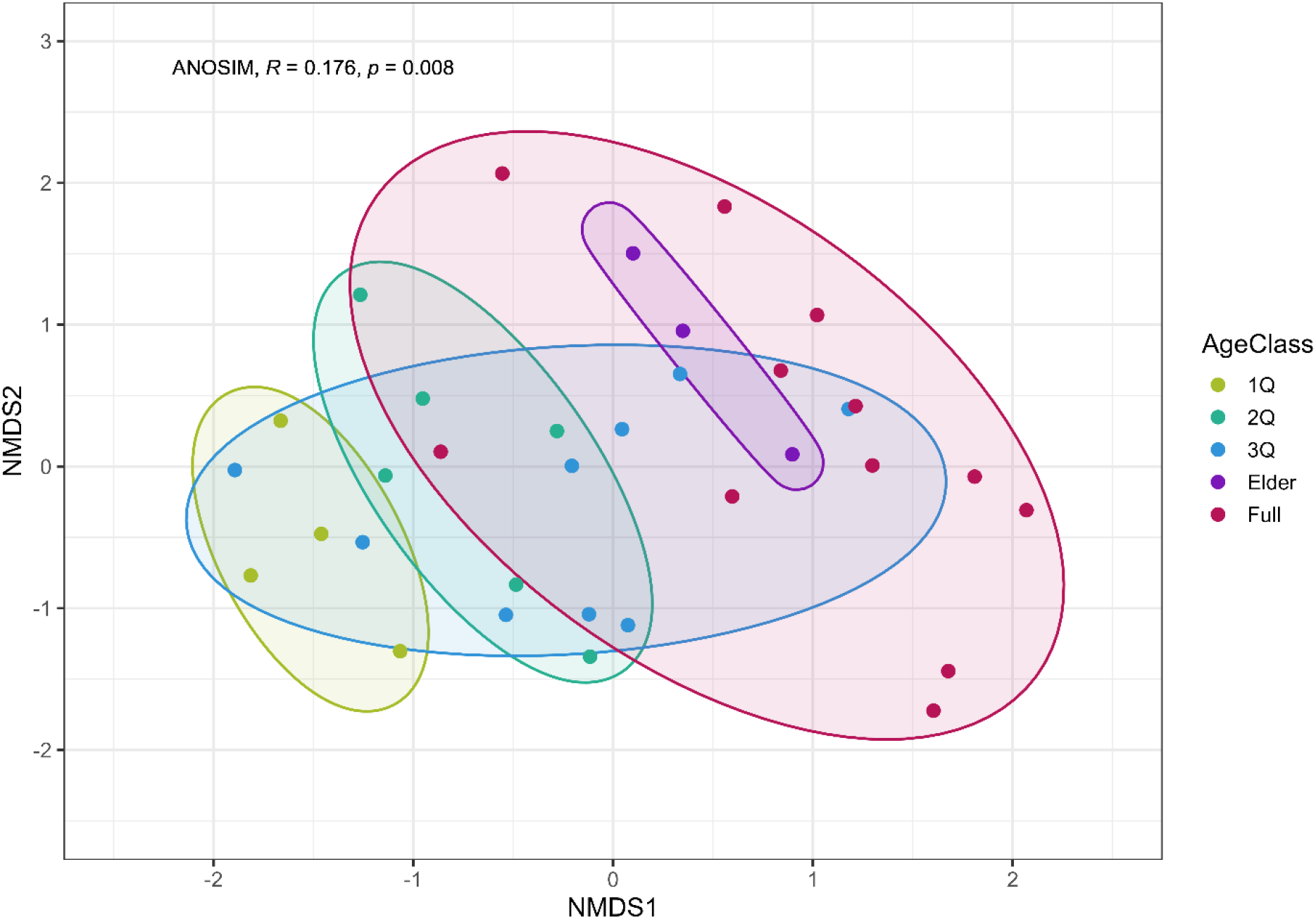
NMDS results when overlaid with the five age classes. Points represent individuals (n = 34), and the colors represent each age class (1Q, n = 4, 2Q, n = 6, 3Q, n = 9, full, n = 12, elder, n = 3). Rather than interpreting where points lie in relation to the axes, the points should be interpreted in relation to each other. Points, or individuals, that are close together have more similarities in character profiles, while individuals farther apart are more dissimilar. ANOSIM analysis revealed significant differences in character profiles between age classes (R = 0.176, *p* = 0.008).

## Discussion

Our study demonstrates both the consistency and flexibility of male elephant behavior across social contexts and time, as well as age-related differences in character profiles. Of the ten behaviors analyzed, affiliation, aggression, dominance, self-directed anxious, and self-directed comfort were significantly repeatable across individuals, while eight of the ten behaviors (i.e., affiliation, aggression, dominance, play, retreat, self-directed anxious, social contentment, and vigilance) were repeatable by event. Vigilance was the only behavioral category that was repeatable by year. The patterns seen in many of these behaviors suggest that individuals consistently express some behaviors differently than others but are also behaviorally adaptable, depending on the social situation.

Keystone and young male presence had the biggest impact on adult males’ behavior rates across the eight repeatable-by-event behaviors; by contrast, social environments with musth males significantly affected only one behavior—retreat. Additionally, we found significant, age-related differences in character profiles for the behaviors that were consistent at the individual level. The youngest age classes (1Q and 2Q) were grouped closely together, occupying a distinctly different space than the full-size and elder individuals. Despite being grouped closely together, many of the individuals displayed idiosyncratic character profiles, particularly in the older age classes (3Q, full, elder), suggesting a large variation in the display of repeatable-by-individual behaviors.

### Behavioral consistency and flexibility

Five behaviors were repeatable-by-individual, which suggests that individual male elephants have distinct character traits (Table 3). These repeatable-by-individual behaviors are those in which individuals behave consistently across time and context, and are also different from each other [2]. As such, our results suggest that male elephants in this population are consistent both in the behaviors they initiate (affiliation, aggression, dominance) and how they respond to the social setting through self-directed behaviors (expressing behaviors that indicate anxiousness or comfort).

The consistent behaviors found in this study reflect both similarities and differences with previous studies of elephant character (or personality) of captive, semi-captive, and free-ranging elephants. Direct comparisons among studies are difficult to make, primarily due to differences in the behaviors assessed, how behaviors were defined, and the sex of the individuals included in the study. However, our results support previous findings of consistency in traits related to sociability and aggression [8–10,13,15–17,21], suggesting the importance of these two traits for elephant character. These two traits are representative of the highly social nature of both males and females, as well as the aggression which occurs within dominance interactions between and within family groups [64–66], and between males [23].

The expression of affiliation and aggression behaviors in males appears to be correlated with position in the social network and dominance hierarchy. Highly socially-integrated males (such as #18, #22, #25, #46, and #65; Figure 1a) display higher and equal proportions of affiliation and aggression behaviors, while the most dominant males (such as #18, #22, #25, #40, and #62; Figure 1b) commonly display higher and equal proportions of affiliation and aggression (#2, #25), or aggression and self-directed anxious (#18, #40, #62) behaviors. This suggests that those that display equal amounts of affiliation and aggression behaviors are the most successful socially, as well as being more dominant. This finding matches our previous study, comparing behaviors across both wet and dry years over a four-year period between 2005-2008, where the most dominant individual (#22) displayed equal amounts of affiliation and aggression behaviors [23]. In our current study, we suggest that individuals who are both highly socially-integrated and dominant balance affiliation and aggression behaviors to maintain bonds as well as their position in the hierarchy. The interplay of dominance and affiliation behaviors is thought to facilitate bonds among individuals within social groups [67] and is the case here as well.

Repeatable-by-event behaviors are those which have rates that are stable within events and differ amongst events. While we expected more flexibility in some behaviors than others in order to cope with a dynamic environment [68], we did not expect four of the five of the repeatable-by-individual behaviors (affiliation, aggression, dominance, and self-direct anxious) to also be repeatable-by-event. Male elephants might have a ‘baseline’ level for each of these four behaviors that are also impacted by other factors, such as the social context or dyadic and group-level relationships. The overlap in individual and event repeatabilities for these four behaviors might be indicative of the ‘individual x environment’ interaction, where individuals are responding differently in the same context [2,69] or have varying levels of behavioral flexibility [68]. This ‘individual x environment’ effect might be important to consider when conducting future elephant character research. Further, repeatability values do not provide insight into the degree of behavioral consistency of individuals, rather an overall population value of the individuals measured [2]. Thus, discerning which repeatable-by-individual behaviors are more susceptible to individual variation and social or environmental context is not possible with repeatability models.

Four behaviors were only repeatable-by-event (play, retreat, social contentment, and vigilance; Table 3). The low individual level repeatability suggests that individuals are more flexible in these behaviors which might be more susceptible to environmental perturbations than the behaviors that were repeatable-by-individual. For example, a behavior such as retreat might be more contextually dependent as it is often a response to aggression behaviors directed towards the focal subject. Similarly, play behaviors often decline with age in male elephants [25,70] and are likely most often initiated by and dependent on the presence of younger individuals [25,71]. For all repeatable behaviors, a large proportion of the variance was unexplained, suggesting several other factors are likely contributing to aspects of behavioral consistency and flexibility in male elephants. Some of these variables might be environmental (e.g. time of day and rainfall conditions), physiological (e.g. hormone concentrations, body condition, and metabolic state), or genetic.

### The influence of social environment on adult male behavior

Given the reputation of musth males being highly aggressive, we expected their presence to have a larger impact on adult male behavior. However, the presence of musth males only had a significant impact on one behavior—retreat (Table 4, Figure 2). When musth and young males are present, the rate of retreat significantly increased for adult males, while just the presence of musth males increased retreat, vigilance, and aggression rates but not significantly. Male elephants rise in the dominance hierarchy when they are in musth [72–74]. Being reproductive competitors, non-musth, age-matched males are likely avoiding aggressive encounters with musth males. Additionally, aggression might be directed only towards individuals who are the highest ranking in the group, rather than blanket aggression towards all males [32]. Musth males might also be more inconsistent in their behavioral expression. In addition, character traits observed outside the state of musth might manifest, perhaps to a lesser degree, during musth. For example, those individuals who are less aggressive and more affiliative outside of musth, might have a different impact on conspecifics during musth than those who are more aggressive. Further research is needed to better understand the nuances of musth male behavior and the impact they have on conspecifics.

As we hypothesized, younger males (age classes 1Q and 2Q) impacted the behavior of adult males (Figure 2, Table 4). Young males are known to solicit and trigger an increase in the rate of affiliation in age-matched and mixed-aged groups [25], which supports our finding. The presence of younger males also significantly increased dominance interactions. Here, and in a previous study [23], adult males were relatively stable in their dominance ranking, while younger males shifted in their position between years. This suggests younger males might be more actively trying to establish themselves in the hierarchy, and older males in the group might be increasing their expression of dominance behaviors to maintain their position, while younger males are present.

Although we expected all social contexts to have an impact on adult male elephant behavior, we were not expecting the presence of both the keystone male and young males to significantly impact the expression of four of the ten behaviors (Figure 2; Table 4). Their presence significantly reduced the expression of aggression, self-directed anxiety, social contentment, and vigilance. The keystone male and his close associates were observed “policing” aggression behavior in other males, particularly younger individuals, an observation that was also noted in another study system [27]. This reduction in aggression might explain the reduced vigilance and self-directed anxiety behaviors observed in the presence of the keystone male, whereby others may have felt less threatened, knowing that such policing would likely occur. Additionally, since the keystone male maintained his position in the dominance hierarchy throughout the study (Figure 1b), the hierarchy might be more structured overall when he is present, further reducing aggression and anxious behaviors related to uncertainty of position in the hierarchy.

Overall, the presence of the keystone male and young males appear to have a positive effect on adult male elephants (reduction of aggression, anxiety, vigilance, and social contentment which implies more social interactions rather than standing around). Older male elephants are important for maintaining social cohesion [24], mediating aggression behaviors [27,75], and functioning as a source of ecological information and effective navigation through the environment [28]. We take these ideas a step further and suggest that particular older individuals, such as the keystone male in our study, and possibly other socially-integrated males, might be an important regulatory element in male elephant society. These results also have implications for *ex-situ* management of male elephant groups, suggesting the need for mixed-age groups and individuals who might take on a more active role in mediating relationships amongst group members. These considerations might also increase the success of the increasingly more common translocations [76]and orphan releases [77,78].

### Age related differences in character profiles

The behavioral profiles of younger adult males (1Q and 2Q) display tight clustering within their age class (Figure 3), indicating that there is less variability in character profiles and personality structures in young males. Males in this age group are transitioning or have recently transitioned into independence and adulthood, exploring away from their family group’s range [79]. On the younger side of this age range, survival is challenging, and only about half of African elephant males survive to 30-35 years old [80]. The tight clustering of behavioral profiles for these age classes suggests limited behavioral strategies to increase chances of survival. Not only are young males seeking out food and water sources, but they also face a drastic change in their social lives, integrating from a predominantly female and family-based life to adult male elephant society [79]. Young males are thought to be forming relationships with peers and older adults to gather environmental knowledge and social skills [25,26,28,79]. These males are also beginning to integrate into local dominance hierarchies, where size and therefore age are important factors in determining position [23]. As such, these males are likely more constrained in how they display these repeatable-by-individual behaviors. However, it remains unclear whether constraint on behavior might create more stability for the duration of the individual’s time in these age classes, or if more flexibility is required to adapt to the changing environment.

After surviving the transition into independence, adult male elephants might have more freedom in their behavioral expression, as indicated by the large variation in character profiles for the 3Q, full, and elder groups (Figure 3). The social niche specialization hypothesis posits that variation in behavior reduces direct competition between individuals [1,81]. For male elephants, intra-specific competition is high, but the adoption of social niches, further facilitated by the maintenance of relatively stable dominance hierarchies [23], might help to reduce conflict. Further, social niches might play a stronger role in species that maintain group membership than species with lower levels of repeated interactions [1]. Our results support this idea, since the individuals included in this study are those who are the most frequent visitors to Mushara, where interactions amongst these individuals are common. Despite apparent social niches in adult male elephants, some individuals are grouped very closely together, suggesting a possible effect of behavioral cohesion amongst individuals with higher association indexes. From our results, age class appears to contribute in large part to driving social niches and possibly behavioral cohesion, but future research is needed to tease apart the impact that age, genetics, association, and dominance have on group dynamics.

### Conclusions

This unique study offers critical insights into the individuality and sociality of free-ranging male elephants. Our results offer a blueprint to establish important aspects of male elephant character, and when combined with further research, may aid in improving conservation policies for male elephants *in-situ* and inform management decisions *ex-situ*. For example, elephant managers in captive settings could consider pairing or grouping appropriate individuals to offer social enrichment based on character traits. *In-situ* managers could use character research to predict the success of translocations or orphan re-integration, and correlate character traits with the propensity to engage in risky foraging behaviors. As human-accelerated climate change, shrinking habitats, poaching, and increased human-elephant conflict continue to put pressure on elephants, understanding the relationship between elephant character, and responses to environmental and social change are critical. Additionally, this research could aid in the development of human-elephant coexistence strategies by accounting for the inter- and intra-individual variation of elephants [82]. Finally, character research can further our understanding of the complexity of male elephant social dynamics, both now and in the future, as they adapt to the changing world, and we can use this knowledge to make more informed conservation management decisions.

## Acknowledgements

The authors thank the Namibian Ministry of Environment and Etosha Ecological Institute for their support of this research. We also thank the contributing volunteers of Utopia Scientific and the Oakland Zoo Conservation Fund for making the field work possible, as well as The Elephant Sanctuary for supporting the data analysis. We thank the Stanford University Vice Provost Office for Undergraduate Education Faculty and Student Grants, and the Smith College Horner Fund Endowment.

## Supporting information

**S1 Table. Overview of the total number of events, intervals, and observation time, as well as the number of individual elephants observed for each year**. Since individual elephants were observed within two or more years, the total elephant number is for unique individuals across the study period.

**S2 Table. Detailed version of the ethogram describing the ten behavioral categories and sixty-six discrete behaviors that make up the behavioral categories.** Behaviors and categories are adapted and modified from [23,37]. For behavior categories, n represents the total occurrence of discrete behaviors within that category, and for discrete behaviors, n represents the total number of times the behavior was displayed. All n values are pooled across individuals, events, and years.

**S3 Table. Results of linear mixed models for each behavior that had a high event repeatability.** Sample sizes (n) are presented as the number of individual elephants (number of behavioral rates). All models included a nested random effect of year, event, and intervals, as well as individual elephant ID as a partially-crossed random effect, with social context as a fixed effect, and the log of the behavior rate as the response variable. Only the behavior rates for non-keystone and non-musth older adults (age classes 3Q, full, and elder) were included in these analyses. The ‘adults’ social context was the reference category for all models, so t and p-values are not provided. Significant social contexts are highlighted with an asterisk. SE = standard error.

## Notes

https://github.com/Utopia-Sci-Lab/Male-Elephant-Character-Durability

### Competing Interest Statement

The authors have declared no competing interest.

### Summary of Updates

Updated the title, added Timothy Rodwell as an author, minor changes to the manuscript, tables, and supporting information.

## References

1. Laskowski KL, Chang C-C, Sheehy K, Aguiñaga J. Consistent Individual Behavioral Variation: What Do We Know and Where Are We Going? Annu Rev Ecol Evol Syst. 2022;53: 161–182. 10.1146/annurev-ecolsys-102220-011451

2. Bell AM, Hankison SJ, Laskowski KL. The repeatability of behaviour: a meta-analysis. Animal Behaviour. 2009;77: 771–783. 10.1016/j.anbehav.2008.12.022

3. Dall SRX, Bell AM, Bolnick DI, Ratnieks FLW. An evolutionary ecology of individual differences. Sih A, editor. Ecol Lett. 2012;15: 1189–1198. doi:10.1111/j.1461-0248.2012.01846.x

4. Carter AJ, Feeney WE, Marshall HH, Cowlishaw G, Heinsohn R. Animal personality: what are behavioural ecologists measuring?: What are animal personality researchers measuring. Biol Rev. 2013;88: 465–475. doi:10.1111/brv.12007

5. MacKinlay RD, Shaw RC. A systematic review of animal personality in conservation science. Conservation Biology. 2022 [cited 18 Nov 2022]. doi:10.1111/cobi.13935

6. Trepel J, Le Roux E, Abraham AJ, Buitenwerf R, Kamp J, Kristensen JA, et al. Meta-analysis shows that wild large herbivores shape ecosystem properties and promote spatial heterogeneity. Nat Ecol Evol. 2024;8: 705–716. 10.1038/s41559-024-02327-6

7. Bergman J, Pedersen RØ, Lundgren EJ, Lemoine RT, Monsarrat S, Pearce EA, et al. Worldwide Late Pleistocene and Early Holocene population declines in extant megafauna are associated with Homo sapiens expansion rather than climate change. Nat Commun. 2023;14: 7679. 10.1038/s41467-023-43426-5

8. Barrett LP, Benson-Amram S. Multiple assessments of personality and problem-solving performance in captive Asian elephants (Elephas maximus) and African savanna elephants (Loxodonta africana). Journal of Comparative Psychology. 2021;135: 406–419. doi:10.1037/com0000281

9. Grand AP, Kuhar CW, Leighty KA, Bettinger TL, Laudenslager ML. Using personality ratings and cortisol to characterize individual differences in African Elephants (*Loxodonta africana*). Applied Animal Behaviour Science. 2012;142: 69–75. doi:10.1016/j.applanim.2012.09.002

10. Horback KM, Miller LJ, Kuczaj SA. Personality assessment in African elephants (*Loxodonta africana*): Comparing the temporal stability of ethological coding versus trait rating. Applied Animal Behaviour Science. 2013;149: 55–62. doi:10.1016/j.applanim.2013.09.009

11. Robertson MR, Olivier LJ, Roberts J, Yonthantham L, Banda C, N’gombwa IB, et al. Testing the Effectiveness of the “Smelly” Elephant Repellent in Controlled Experiments in Semi-Captive Asian and African Savanna Elephants. Animals. 2023;13: 3334. doi:10.3390/ani13213334

12. Rutherford L, Murray LE. Personality and behavioral changes in Asian elephants (*Elephas maximus*) following the death of herd members. Integr Zool. 2020;16: 170–188. doi:10.1111/1749-4877.12476

13. Williams E, Carter A, Hall C, Bremner-Harrison S. Exploring the relationship between personality and social interactions in zoo-housed elephants: Incorporation of keeper expertise. Applied Animal Behaviour Science. 2019;221: 104876. doi:10.1016/j.applanim.2019.104876

14. Williams E, Carter A, Hall C, Bremner-Harrison S. Social Interactions in Zoo-Housed Elephants: Factors Affecting Social Relationships. Animals. 2019;9: 747. doi:10.3390/ani9100747

15. Yasui S, Konno A, Tanaka M, Idani G, Ludwig A, Lieckfeldt D, et al. Personality Assessment and Its Association With Genetic Factors in Captive Asian and African Elephants: Elephant Personality and Genetic Factors. Zoo Biol. 2013;32: 70–78. doi:10.1002/zoo.21045

16. Seltmann MW, Helle S, Adams MJ, Mar KU, Lahdenperä M. Evaluating the personality structure of semi-captive Asian elephants living in their natural habitat. R Soc open sci. 2018;5: 172026. doi:10.1098/rsos.172026

17. Seltmann MW, Helle S, Htut W, Lahdenperä M. Males have more aggressive and less sociable personalities than females in semi-captive Asian elephants. Sci Rep. 2019;9: 2668. doi:10.1038/s41598-019-39915-7

18. Webb JL, Crawley JAH, Seltmann MW, Liehrmann O, Hemmings N, Nyein UK, et al. Evaluating the Reliability of Non-Specialist Observers in the Behavioural Assessment of Semi-Captive Asian Elephant Welfare. Animals. 2020;10: 167. 10.3390/ani10010167

19. Srinivasaiah NM, Varma S, Sukumar R. Documenting Indigenous Traditional Knowledge of the Asian Elephant in Captivity. Bangalore 560012, India: Asian Nature Conservation Foundation (ANCF), c/o Centre for Ecological Sciences, Indian Institute of Science; 2014. Available: http://www.asiannature.org/sites/default/files/Elephants%20and%20Mahouts-%20Final%20Report%205March14.pdf

20. Beirne C, Houslay TM, Morkel P, Clark CJ, Fay M, Okouyi J, et al. African forest elephant movements depend on time scale and individual behavior. Sci Rep. 2021;11: 12634. 10.1038/s41598-021-91627-z

21. Lee PC, Moss CJ. Wild female African elephants (*Loxodonta africana*) exhibit personality traits of leadership and social integration. Journal of Comparative Psychology. 2012;126: 224–232. doi:10.1037/a0026566

22. Moss CJ, Poole JH. Relationships and social structure in African elephants. In: Hinde RA, editor. Primate Social Relationships: an Integrated Approach. Oxford: Blackwell Scientific Publications; 1983.

23. O’Connell-Rodwell CE, Wood JD, Kinzley C, Rodwell TC, Alarcon C, Wasser SK, et al. Male African elephants (*Loxodonta africana*) queue when the stakes are high. Ethology Ecology & Evolution. 2011;23: 388–397. 10.1080/03949370.2011.598569

24. Chiyo PI, Archie EA, Hollister-Smith JA, Lee PC, Poole JH, Moss CJ, et al. Association patterns of African elephants in all-male groups: the role of age and genetic relatedness. Animal Behaviour. 2011;81: 1093–1099. 10.1016/j.anbehav.2011.02.013

25. Evans KE, Harris S. Adolescence in male African elephants, *Loxodonta africana*, and the importance of sociality. Animal Behaviour. 2008;76: 779–787. 10.1016/j.anbehav.2008.03.019

26. Murphy D, Mumby HS, Henley MD. Age differences in the temporal stability of a male African elephant (*Loxodonta africana*) social network. Griffin A, editor. Behavioral Ecology. 2019; arz152. doi:10.1093/beheco/arz152

27. Allen CRB, Croft DP, Brent LJN. Reduced older male presence linked to increased rates of aggression to non-conspecific targets in male elephant. Proceedings of the Royal Society B. 2021;288: 20211374. 10.1098/rspb.2021.1374

28. Allen CRB, Brent LJN, Motsentwa T, Weiss MN, Croft DP. Importance of old bulls: leaders and followers in collective movements of all-male groups in African savannah elephants (Loxodonta africana). Sci Rep. 2020;10: 13996. 10.1038/s41598-020-70682-y

29. Thouless CR, Dublin HT, Blanc JJ, Skinner DP, Daniels TE, Taylor RD, et al. African elephant status report 2016: an update from the African elephant database. Gland, Switzerland: IUCN/ SSC African Elephant Specialist Group; 2016. Report No.: 60. doi:10.2305/IUCN.CH.2007.SSC-OP.33.en

30. Craig GC, St. C Gibson D, Uiseb KH. Namibia’s elephants—population, distribution and trends. Pachyderm. 2021;62: 35–52. doi:https://pachydermjournal.org/index.php/pachyderm/article/view/460

31. Thurber MI, O’Connell-Rodwell CE, Turner WC, Nambandi K, Kinzley C, Rodwell TC, et al. Effects of rainfall, host demography, and musth on strongyle fecal egg counts in African elephants (*Loxodonta africana*) in Namibia. Journal of Wildlife Diseases. 2011;47: 172–181. 10.7589/0090-3558-47.1.172

32. O’Connell-Rodwell CE, Sandri MN, Berezin JL, Munevar JM, Kinzley C, Wood JD, et al. Male African Elephant (Loxodonta africana) Behavioral Responses to Estrous Call Playbacks May Inform Conservation Management Tools. Animals. 2022;12: 1162. 10.3390/ani12091162

33. Berezin JL, Odom AJ, Hayssen V, O’Connell-Rodwell CE. A Snapshot into the Lives of Elephants: Camera Traps and Conservation in Etosha National Park, Namibia. Diversity. 2023;15: 1146. 10.3390/d15111146

34. Moss C. Getting to know a Population. In: Kangwana K, editor. Studying Elephants. Nairobi, Kenya: African Wildlife Foundation; 1996. pp. 58–74.

35. O’Connell-Rodwell CE, Freeman PT, Kinzley C, Sandri MN, Berezin JL, Wiśniewska M, et al. A novel technique for aging male African elephants (*Loxodonta africana*) using craniofacial photogrammetry and geometric morphometrics. Mamm Biol. 2022;102: 591–613. 10.1007/s42991-022-00238-2

36. Altmann J. Observational Study of Behavior: Sampling Methods. Behav. 1974;49: 227–266. doi:10.1163/156853974X00534

37. Poole J, Granli P. The Elephant Ethogram: a library of African elephant behaviour. Pachyderm. 2021;62: 105–111. Available: https://pachydermjournal.org/index.php/pachyderm/article/view/462

38. Franks DW, Ruxton GD, James R. Sampling animal association networks with the gambit of the group. Behav Ecol Sociobiol. 2010;64: 493–503. 10.1007/s00265-009-0865-8

39. Cairns SJ, Schwager SJ. A comparison of association indices. Animal Behaviour. 1987;35: 1454–1469. 10.1016/S0003-3472(87)80018-0

40. Whitehead H. Analyzing Animal Societies: Quantitative Methods for Vertebrate Social Analysis. University of Chicago Press; 2008. Available: https://press.uchicago.edu/ucp/books/book/chicago/A/bo5607202.html

41. Wasserman S, Faust K. Social Network Analysis: Methods and Applications. 1st ed. Cambridge University Press; 1994. doi:10.1017/CBO9780511815478

42. Farine DR. Animal social network inference and permutations for ecologists in R using *asnipe*. Methods Ecol Evol. 2013;4: 1187–1194. 10.1111/2041-210X.12121

43. Csárdi G, Nepusz T. The Igraph Software Package for Complex Network Research. InterJournal, complex systems. 2006;1695: 1–9. Available: http://igraph.org

44. David HA. Ranking from unbalanced paired-comparison data. Biometrika. 1987;74: 432–436. 10.1093/biomet/74.2.432

45. de Vries H, Stevens JMG, Vervaecke H. Measuring and testing the steepness of dominance hierarchies. Animal Behaviour. 2006;71: 585–592. 10.1016/j.anbehav.2005.05.015

46. Gammell MP, De Vries H, Jennings DJ, Carlin CM, Hayden TJ. David’s score: a more appropriate dominance ranking method than Clutton-Brock et al.’s index. Animal Behaviour. 2003;66: 601– 605. 10.1006/anbe.2003.2226

47. Fujii K, Jin J, Vandeleest J, Shev A, Beisner B, McCowan B, et al. Perc: Using Percolation and Conductance to Find Information Flow Certainty in a Direct Network. 2021. Available: https://CRAN.R-project.org/package=Perc.

48. Neumann C, Kulik L. EloRating: Animal Dominance Hierarchies by Elo Rating. 2020. Available: https://CRAN.R-project.org/package=EloRating

49. R Core Team. R: A language and environment for statistical computing. Vienna, Austria: R Foundation for Statistical Computing; 2023. Available: https://www.R-project.org/

50. Stoffel MA, Nakagawa S, Schielzeth H. rptR: repeatability estimation and variance decomposition by generalized linear mixed-effects models. Goslee S, editor. Methods Ecol Evol. 2017;8: 1639–1644. 10.1111/2041-210X.12797

51. Nakagawa S, Schielzeth H. Repeatability for Gaussian and non-Gaussian data: a practical guide for biologists. Biological Reviews. 2010; no-no. 10.1111/j.1469-185X.2010.00141.x

52. Schuster AC, Carl T, Foerster K. Repeatability and consistency of individual behaviour in juvenile and adult Eurasian harvest mice. Sci Nat. 2017;104: 10. 10.1007%2Fs00114-017-1430-3

53. Kuznetsova A, Brockhoff PB, Christensen RHB. lmerTest Package: Tests in Linear Mixed Effects Models. J Stat Soft. 2017;82. 10.18637/jss.v082.i13

54. Daniel Lüdecke. ggeffects: Tidy Data Frames of Marginal Effects from Regression Models. Journal of Open Source Software. 2018;3: 772. doi:10.21105/joss.00772

55. Florian Hartig. DHARMa: Residual Diagnostics for Hierarchical (Multi-Level / Mixed) Regression Models. 2022. Available: https://CRAN.R-project.org/package=DHARMa

56. Budaev SV. Using Principal Components and Factor Analysis in Animal Behaviour Research: Caveats and Guidelines. Ethology. 2010;116: 472–480. doi:10.1111/j.1439-0310.2010.01758.x

57. Quinn GP, Keough MJ. Experimental design and data analysis for biologists. Cambridge: Cambridge University Press; 2002.

58. Aitchison J. The Statistical Analysis of Compositional Data. Journal of the Royal Statistical Society: Series B (Methodological). 1982;44: 139–160. 10.1111/j.2517-6161.1982.tb01195.x

59. Aitchison J, Barceló-Vidal C, Martín-Fernández JA, Pawlowsky-Glahn V. Logratio Analysis and Compositional Distance. Mathematical Geology. 2000;32: 271–275. 10.1023/A:1007529726302

60. Quinn TP, Erb I, Richardson MF, Crowley TM. Understanding sequencing data as compositions: an outlook and review. Wren J, editor. Bioinformatics. 2018;34: 2870–2878. 10.1093/bioinformatics/bty175

61. Martino C, Morton JT, Marotz CA, Thompson LR, Tripathi A, Knight R, et al. A Novel Sparse Compositional Technique Reveals Microbial Perturbations. Neufeld JD, editor. mSystems. 2019;4: e00016–19. 10.1128/mSystems.00016-19

62. Oksanen J, Simpson G, Blanchet F, Kindt R, Legendre P, Minchin P, et al. vegan: Community Ecology Package. 2022. Available: https://CRAN.R-project.org/package=vegan

63. Clarke KR. Non-parametric multivariate analyses of changes in community structure. Austral Ecol. 1993;18: 117–143. 10.1111/j.1442-9993.1993.tb00438.x

64. Wittemyer G, Polansky L, Douglas-Hamilton I, Getz WM. Disentangling the effects of forage, social rank, and risk on movement autocorrelation of elephants using Fourier and wavelet analyses. Proc Natl Acad Sci USA. 2008;105: 19108–19113. 10.1073/pnas.0801744105

65. Wittemyer G, Getz WM, Vollrath F, Douglas-Hamilton I. Social dominance, seasonal movements, and spatial segregation in African elephants: a contribution to conservation behavior. Behav Ecol Sociobiol. 2007;61: 1919–1931. 10.1007/s00265-007-0432-0

66. Wittemyer G, Getz WM. Hierarchical dominance structure and social organization in African elephants, Loxodonta africana. Animal Behaviour. 2007;73: 671–681. 10.1016/j.anbehav.2006.10.008

67. de Waal FBM. The Integration of Dominance and Social Bonding in Primates. The Quarterly Review of Biology. 1986;61: 459–479. 10.1086/415144

68. Dingemanse NJ, Kazem AJN, Réale D, Wright J. Behavioural reaction norms: animal personality meets individual plasticity. Trends in Ecology & Evolution. 2010;25: 81–89. 10.1016/j.tree.2009.07.013

69. Martin JGA, Réale D. Temperament, risk assessment and habituation to novelty in eastern chipmunks, Tamias striatus. Animal Behaviour. 2008;75: 309–318. 10.1016/j.anbehav.2007.05.026

70. Lee PC, Moss CJ. Calf Development and Maternal Rearing Strategies. In: Moss CJ, Croze H, Lee PC, editors. The Amboseli Elephants: A Long-Term Perspective on a Long-Lived Mammal. Chicago: University of Chicago Press; 2011. pp. 224–237.

71. Freeman PT, Anderson EL, Allen KB, O’Connell-Rodwell CE. Age-based variation in calf independence, social behavior and play in a captive population of African elephant calves. Zoo Biology. 2021;40: 376–385. doi:10.1002/zoo.21629

72. Chelliah K, Sukumar R. The role of tusks, musth and body size in male–male competition among Asian elephants, *Elephas maximus*. Animal Behaviour. 2013;86: 1207–1214. 10.1016/j.anbehav.2013.09.022

73. Hollister-Smith JA, Poole JH, Archie EA, Vance EA, Georgiadis NJ, Moss CJ, et al. Age, musth and paternity success in wild male African elephants, *Loxodonta africana*. Animal Behaviour. 2007;74: 287–296. 10.1016/j.anbehav.2006.12.008

74. Poole JH. Announcing intent: the aggressive state of musth in African elephants. Animal Behaviour. 1989;37: 140–152. 10.1016/0003-3472(89)90014-6

75. Slotow R, van Dyk G, Poole J, Page B, Klocke A. Older bull elephants control young males. Nature. 2000;408: 425–426. 10.1038/35044191

76. Tiller LN, King LE, Okita-Ouma B, Lala F, Pope F, Douglas-Hamilton I, et al. The behaviour and fate of translocated bull African savanna elephants (*Loxodonta africana*) into a novel environment. Afr J Ecol. 2022; aje.13038. doi:10.1111/aje.13038

77. Goldenberg SZ, Chege SM, Mwangi N, Craig I, Daballen D, Douglas-Hamilton I, et al. Social integration of translocated wildlife: a case study of rehabilitated and released elephant calves in northern Kenya. Mamm Biol. 2022;102: 1299–1314. 10.1007/s42991-022-00285-9

78. Greggor AL, Blumstein DT, Wong BBM, Berger-Tal O. Using animal behavior in conservation management: a series of systematic reviews and maps. Environ Evid. 2019;8: 23, s13750–019-0164–4. doi:10.1186/s13750-019-0164-4

79. Lee PC, Poole JH, Njiraini, Norah, Sayialel, Catherine N., Moss, Cynthia J. Chapter 17: Male social dynamics: independence and beyond. In: Moss CJ, Croze H, Lee PC, editors. The Amboseli Elephants: A Long-Term Perspective on a Long-Lived Mammal. Chicago: University of Chicago Press; 2011. pp. 260–271. Available: 10.7208/chicago/9780226542263.001.0001

80. Moss CJ. The demography of an African elephant (*Loxodonta africana*) population in Amboseli, Kenya. Journal of Zoology. 2001;255: 145–156. doi:10.1017/S0952836901001212

81. Bergmüller R, Taborsky M. Animal personality due to social niche specialisation. Trends in Ecology & Evolution. 2010;25: 504–511. doi:10.1016/j.tree.2010.06.012

82. Mumby HS, Plotnik JM. Taking the Elephants’ Perspective: Remembering Elephant Behavior, Cognition and Ecology in Human-Elephant Conflict Mitigation. Front Ecol Evol. 2018;6: 122. 10.3389/fevo.2018.00122

