## Supplementary material for "Dynamics of male African elephant character durability across time, age, and social contexts": S1 Table

| **Year** | **Events** | **Intervals** | **Total observation time (minutes)** | **Elephants (n)** |
| --- | --- | --- | --- | --- |
| 2007 | 37 | 47 | 3,124.5 | 23 |
| 2008 | 17 | 25 | 1,186.4 | 14 |
| 2009 | 40 | 59 | 2,794.6 | 28 |
| 2010 | 34 | 41 | 2,310.4 | 23 |
| 2011 | 20 | 28 | 964.6 | 18 |
| *Total* | *148* | *200* | *10,380.5* | *34* |
