## Supplementary material for "Dynamics of male African elephant character durability across time, age, and social contexts": S2 Table

| **Behavior**  **(category)** | **Behavior category**  **definition** | **Behavior**  **(fine-scale)** | **n** | **Behavior description** |
| --- | --- | --- | --- | --- |
| Affiliation  (n = 2,461) | Positive interaction between individuals; aids in relationship building and maintenance. | Backs into | 3 | Elephant gently backs into another, resulting in body contact. |
|  |  | Body to body | 160 | Elephant touches the body of another with his body. |
|  |  | Ear on face | 34 | Elephant places his ear over the head or face of another. |
|  |  | Ear on rear | 48 | Elephant places his ear over the rear of another. |
|  |  | Follows | 11 | Elephant follows behind, in the same direction as a conspecific. |
|  |  | Foot to body | 1 | Elephant touches the body of another, using his front or hind foot. |
|  |  | Head to body | 250 | Elephant touches the body of another with his head. |
|  |  | Head to head | 502 | Elephant touches the head of another elephant with his head. |
|  |  | Inspecting | 78 | One elephant reaches his trunk towards another from a body length or farther than one body length away. |
|  |  | Mount | 1 | Elephant lifts his front legs and stands on the back of another elephant and attempts penetration. |
|  |  | Premount | 66 | Elephant positions his head on top of another elephant’s hindquarters and rests his trunk along the spine of the other. May or may not result in a mount. |
|  |  | Pushes | 113 | Elephant pushes another in an affiliative fashion, usually upon leaving the waterhole. |
|  |  | Rubs | 7 | Elephant scratches against a body part of another. |
|  |  | Tail to body | 101 | Elephant touches anywhere on the body of another using his tail. |
|  |  | Trunk to body | 20 | Elephant uses his trunk to touch the body of another (excluding head or temporal region). |
|  |  | Trunk to head | 271 | Elephant touches the head of another elephant with his trunk. |
|  |  | Trunk to mouth | 703 | Elephant touches and places his trunk in the mouth of another. |
|  |  | Trunk to temporal | 39 | Elephant touches the temporal region of another, using his trunk. |
|  |  | Trunk to tusk | 9 | Elephant uses his trunk to touch the tusk of another. |
|  |  | Trunk wrap | 44 | Two elephants intertwine trunks. |
| Aggression  (n = 2,038) | Negative interaction to intimidate or threaten other elephants; no physical contact made between elephants. | Aggressive ear flap | 57 | Flaps both ears forward and backward aggressively in threat, often with head held high. |
|  |  | Ear fold | 65 | Bending ears backward just below the midline, may be associated with “ears held out” and “head held up”. |
|  |  | Ears held out | 1,278 | Aggressive threat – both ears rigidly held in an extended position, often with “head held up.” |
|  |  | Foot toss | 294 | Distant threat – one front foot is tosses aggressively, or swung in the direction of the recipient, kicking the air. |
|  |  | Hard ear flap | 2 | Slaps ears aggressively against body, usually making a sound. Often observed with aggressive ear flap. |
|  |  | Head held up | 62 | Head raised higher than the shoulders, usually with “ears held out.” |
|  |  | Head shake | 72 | Abrupt shaking of the head, causing ears to flap, making a “cracking” sound. |
|  |  | Open mouth threat | 91 | Mouth held open for an extended period, often accompanied with “head held up”, brief “head down”, or with an “ear fold” as the aggressor approaches another. This might also be used when in retreat. |
|  |  | Tail slap | 17 | Elephant slaps his tail against his own backside, usually in an up and down motion, facing the recipient. |
|  |  | Trunk drag | 90 | Elephant walks slowly with a length of trunk contacting the ground, often making a rough sound against the earth. |
|  |  | Trunk fist | 1 | Trunk tightly balled up into a fist shape, often precedes trunk throw. |
|  |  | Trunk throw | 9 | Trunk swung out towards recipient. |
| Dominance  (n = 872) | Displacement of another individual; used to calculate the dominance hierarchy. | Displacement | 872 | One elephant forces another to change his position and move away, possibly so the initiator can occupy the position or to exert dominance; displacement occurs with body contact or without. |
| Escalated  aggression  (n = 69) | Negative interaction to threaten or attach other elephants; often no physical contact. | Charge | 5 | One elephant rushes towards another, usually with head held up and ears held out, or folded; might stop short of recipient and include a “trunk throw,” “foot toss”; may be associated with a vocalization from the recipient if particularly intense. |
|  |  | Chase | 14 | Persistent, prolonged, and aggressive follow, in a fast walk pace. |
|  |  | Combat | 6 | Aggressive contact between two elephants; rushing towards each other with trunks curled under to increase tusk contact; might lower head on approach. |
|  |  | Head down | 2 | Single slight dipping of the head, usually while rushing toward recipient, sometimes combined with ear fold or ears held out. |
|  |  | Head thrust | 32 | Abrupt throwing of the head forward towards the recipient, while holding head high. |
|  |  | Lunge | 3 | Abrupt forward step with head thrust towards the recipient, with head held high. |
|  |  | Pursue | 2 | Aggressive follow at a walking pace. |
|  |  | Stand off | 5 | Two males stand facing each other, possibly preceding and/or during combat. |
| Play  (n = 493) | Social play where individuals engage in exaggerated or loose movements and friendly sparring that avoids injury. | Gentle sparing | 484 | Two elephants, usually in head-to-head position, pushing back and forth, usually using tusks and trunk, might evolve into a rougher encounter. |
|  |  | Trunk swing | 9 | One elephant loosely swings his trunk from side to side or front to back in an affiliative context. |
| Retreat  (n = 206) | Avoidance of aggression from another individual. | Back up | 100 | Without turning away from another individual, elephant slowly backs away in the opposite direction. |
|  |  | Retreat | 106 | Elephant turns and moves quickly in the opposite direction of a perceived threat. |
| Self-directed anxious  (n = 1,882) | Nervous and reactive behaviors directed towards oneself; often to attend to the actions of other elephants or after receiving aggression from another. | Foot rubs | 89 | Rubbing front or back foot on the other, sometimes when unsure of what action to take. |
|  |  | Tail lift | 9 | Elephant lifting tail in retreat from a potential threat. |
|  |  | Touches own tusk | 158 | Trunk tip touches own tusk, usually in “dabbing” motion with repetitive touching, sometimes also includes grabbing and pulling own tusk. |
|  |  | Trunk own mouth | 728 | Trunk tip touches own mouth, not for water or flehmen behavior, but in apparent uncertainty about the social situation. |
|  |  | Trunk own temporal | 75 | Trunk tip touches own temporal region on the head. |
|  |  | Trunk suck | 573 | Trunk tip placed in own mouth while sucking in, not associated with drinking. |
|  |  | Trunk twist | 250 | Trunk twists and untwists at a moderate or fast pace. |
| Self-directed comfort  (n = 498) | Comfort or relaxed behaviors directed towards oneself. | Cross leg | 1 | Standing with back legs crossed, one over the other. |
|  |  | Rest trunk | 23 | Resting or standing elephant with fully-relaxed trunk laying flaccid on the ground. Distinct from listening behavior, which includes a combination of behaviors with ears held out at 45-degree angle, alert posture, and more rigid trunk. |
|  |  | Tusk hang | 474 | Elephant drapes trunk over his own tusk. Trunk is usually flaccid/relaxed. |
| Social  Contentment  (n = 1,992) | Group activity; relaxed behaviors observed during times of stillness at the waterhole. | Ear flap | 1,078 | Elephants flap their ears and swing their tails back and forth (like a pendulum), directing their gaze at each other; usually initiated when they’re standing still. |
|  |  | Tailswing | 914 | Elephants flap their ears and swing their tails back and forth (like a pendulum), directing their gaze at each other; usually initiated when they’re standing still. |
| Vigilance  (n = 4,616) | Environmental assessment, often towards a conspecific to attend to the actions of other elephants; vigilance behaviors directed toward human activity were removed. | Freeze | 3 | Elephant immediately stops all movement. |
|  |  | Freeze trunk ground | 9 | Elephant immediately stops all movement and places his trunk on the ground rigidly, possibly to listen or detect movement. |
|  |  | Look | 1,133 | An elephant is alert and intently facing a specific direction or conspecific or object, with an obvious opening of the eyes. |
|  |  | Orient | 328 | Elephant turns his entire body towards another elephant. |
|  |  | Over shoulder | 2,123 | Without orienting his body in the direction of another, an elephant looks over his shoulder at a conspecific (either visually present or in the distance). |
|  |  | Smell | 1,020 | Elephant uses “periscope trunk,” where trunk is extended high above the head in the direction of a conspecific or some novel item or disturbance (either visually present or in the distance). |
