## Supplementary material for "Dynamics of male African elephant character durability across time, age, and social contexts": S3 Table

| **Behavior** | **n** | **Social context** | **Estimate** | **SE** | ***t*-statistic** | ***p*-value** |
| --- | --- | --- | --- | --- | --- | --- |
| Affiliation | 23(265) | Adults | -2.50 | 0.14 |  |  |
|  |  | Keystone | -0.72 | 0.27 | -2.69 | 0.009 * |
|  |  | Keystone & Youth | -0.12 | 0.19 | -0.57 | 0.574 |
|  |  | Musth | -0.48 | 0.27 | -1.77 | 0.079 |
|  |  | Musth & Youth | -0.04 | -0.04 | 0.32 | 0.909 |
|  |  | Youth | 0.41 | 0.18 | 2.34 | 0.021 * |
| Aggression | 23(318) | Adults | -2.53 | 0.14 |  |  |
|  |  | Keystone | -0.11 | 0.23 | -0.47 | 0.643 |
|  |  | Keystone & Youth | -0.46 | 0.18 | -2.49 | 0.014 * |
|  |  | Musth | 0.32 | 0.21 | 1.56 | 0.121 |
|  |  | Musth & Youth | 0.24 | 0.28 | 0.85 | 0.398 |
|  |  | Youth | 0.10 | 0.16 | 0.64 | 0.525 |
| Dominance | 23(216) | Adults | -3.27 | 0.14 |  |  |
|  |  | Keystone | 0.27 | 0.24 | 1.13 | 0.263 |
|  |  | Keystone & Youth | 0.34 | 0.18 | 1.90 | 0.062 |
|  |  | Musth | 0.27 | 0.28 | 0.98 | 0.328 |
|  |  | Musth & Youth | 0.51 | 0.28 | 1.83 | 0.069 |
|  |  | Youth | 0.58 | 0.16 | 3.52 | < 0.001 * |
| Play | 20(87) | Adults | -2.93 | 0.34 |  |  |
|  |  | Keystone | -0.40 | 0.55 | -0.73 | 0.472 |
|  |  | Keystone & Youth | -0.29 | 0.42 | -0.69 | 0.492 |
|  |  | Musth | 0.22 | 0.67 | 0.33 | 0.746 |
|  |  | Musth & Youth | -0.75 | 0.83 | -0.91 | 0.365 |
|  |  | Youth | 0.61 | 0.36 | 1.69 | 0.098 |
| Retreat | 21(75) | Adults | -3.41 | 0.21 |  |  |
|  |  | Keystone | -0.25 | 0.57 | -0.45 | 0.656 |
|  |  | Keystone & Youth | -0.42 | 0.35 | -1.19 | 0.241 |
|  |  | Musth | 0.67 | 0.36 | 1.86 | 0.070 |
|  |  | Musth & Youth | 0.82 | 0.41 | 2.01 | 0.049 * |
|  |  | Youth | 0.30 | 0.44 | 0.69 | 0.492 |
| Self-directed anxious | 23(293) | Adults | -2.62 | 0.21 |  |  |
|  |  | Keystone | -0.32 | 0.21 | -1.58 | 0.118 |
|  |  | Keystone & Youth | -0.50 | 0.16 | -3.06 | 0.003 * |
|  |  | Musth | -0.02 | 0.20 | -0.08 | 0.938 |
|  |  | Musth & Youth | 0.21 | 0.31 | 0.68 | 0.496 |
|  |  | Youth | -0.05 | 0.15 | -0.36 | 0.717 |
| Social contentment | 23(261) | Adults | -2.30 | 0.19 |  |  |
|  |  | Keystone | -0.15 | 0.26 | -0.57 | 0.568 |
|  |  | Keystone & Youth | -0.73 | 0.22 | -3.28 | 0.001 * |
|  |  | Musth | -0.25 | 0.27 | -0.90 | 0.371 |
|  |  | Musth & Youth | -0.68 | 0.36 | -1.86 | 0.065 |
|  |  | Youth | -0.31 | 0.19 | -1.58 | 0.117 |
| Vigilance | 23(383) | Adults | -1.58 | 0.16 |  |  |
|  |  | Keystone | -0.35 | 0.19 | -1.83 | 0.070 |
|  |  | Keystone & Youth | -0.70 | 0.16 | -4.48 | < 0.001 * |
|  |  | Musth | 0.29 | 0.17 | 1.67 | 0.096 |
|  |  | Musth & Youth | 0.11 | 0.22 | 0.50 | 0.620 |
|  |  | Youth | -0.31 | 0.13 | -2.34 | 0.021 * |
